# Bibliome Variant Database: Automated Identification and Annotation of Genetic Variants in Primary Literature

**DOI:** 10.1101/2020.07.16.207688

**Authors:** Samuel W. Baker, Arupa Ganguly

## Abstract

The Bibliome Variant Database (BVdb) is a freely available reference database containing over 1 million human genetic variants mapped to the human genome that have been mined from primary literature. The BVdb is designed to facilitate variant interpretation in clinical and research contexts by reducing or eliminating the time required to search for literature describing a given variant. Users can search the database using gene symbols, HGVS variant nomenclature, genomic positions, or rsIDs. Each variant page lists references in the database that describe the variant, as well as the exact gene symbol and variant text description identified in each reference.

**AVAILABILITY AND IMPLEMENTATION:** The BVdb is freely available at http://bibliome.ai

## INTRODUCTION

Clinical exome and genome sequencing are increasingly being used to diagnose rare genetic disease and in the treatment of cancer (Williams 2019). Such methods identify hundreds of genomic variants that must be evaluated to determine which are most relevant to a patient’s disease (Retterer et al. 2016). In the majority of clinical laboratories, variant interpretation is a labor-intensive manual process and interpretation of each variant can require up to one hour or more of expert time (Dewey et al. 2014). Literature review plays a central role in variant interpretation and information gathered from primary sources describing a given variant is routinely used during application of germline and somatic variant classification guidelines (Richards et al. 2015; Li et al. 2017). Curated variant interpretation databases, such as ClinVar, Human Gene Mutation Database (HGMD), and Clinical Interpretation of Variants in Cancer (CIVIC), represent valuable resources linking genomic variants to primary literature, and their use enables more efficient variant interpretation by reducing or eliminating the time required to search for primary literature describing a given variant (Landrum et al. 2016; Stenson et al. 2017; Griffith et al. 2017). However, owing to the significant labor requirements of manual curation, such databases include only a proportion of all published variants (Stenson et al. 2017; Griffith et al. 2017).

Here, we present the Bibliome Variant Database (BVdb), a structured variant reference database generated through automated identification and annotation of genomic variants in open access literature. The BVdb is designed to facilitate variant interpretation in clinical, diagnostic, and research contexts by allowing rapid lookup of primary literature describing genetic variation across genes and genomic loci.

## MATERIALS AND METHODS

The BVdb consists of a text mining pipeline and website, both of which were written in Python. The PubMed Central Open Access Subset and Author Manuscript Collection, together containing over 3.6 million full-text publications, was downloaded from the NCBI FTP server on June 10^th^, 2020 (ftp://ftp.ncbi.nlm.nih.gov/pub/pmc/). Additional open access publications referenced in the Unpaywall Database Snapshot were also downloaded (https://unpaywall.org/products/snapshot, last accessed June 10th, 2020). For each publication, regular expressions were used to identify rsIDs as well as text patterns corresponding to HGVS-compliant and commonly used HGVS-non-compliant variant nomenclature describing cDNA and protein sequence changes (den Dunnen et al. 2016). Each variant identified within a given publication was assigned to a gene symbol present in the same publication based on spatial, numeric, and contextual features within the text. Gene-assigned variants were added to the BVdb only if the cDNA and/or protein annotations could be mapped to both the canonical transcript, or transcript present in the text, of the assigned gene and the human genome version hg19. For incomplete variant annotations, i.e. cDNA annotations identified in the absence of a corresponding protein annotation and vise-versa, the absent cDNA or protein annotation was imputed and the combination of the identified and imputed variant annotations were added to the BVdb. Genomic coordinates for each gene-assigned variant were determined using the Transvar reverse annotation function, and variant annotations displayed in the BVdb browser were generated using VEP (Zhou et al. 2015; McLaren et al. 2016).

## RESULTS AND DISCUSSION

The BVdb can be accessed at https://bibliome.ai using a desktop computer or mobile device. Contents of the BVdb are presented in three basic interconnected views, each of which is designed to provide information content in a format which facilitates variant interpretation and classification.

1. The gene view (Figure 1A) displays all variants in the BVdb assigned to a given gene and can be accessed by entering an HGNC-approved gene symbol in the search bar. Gene-specific information, including transcript used for annotation and links to external gene-specific resources, is located at the top of the page. Variants listed in the gene view are sorted by genomic position and information including cDNA annotation, protein annotation, predicted coding effect, exon/intron number, and number of references in which the variant was identified is displayed for each variant.
2. The variant view (Figure 1B) can be accessed either by clicking on a variant ID on the gene page or by entering a variant-specific search term in the search bar. Each variant page lists publications in which the variant was identified and provides links to variant-specific entries in external databases. Each publication entry contains information including, article title, author list, journal, publication date, and the specific variant representation identified in the article text (i.e. *KRAS* p.Gly12Asp versus *KRAS* p.G12D).
3. The publication view (Figure 1C) can be accessed by clicking on PMIDs in the variant view or by entering a PMID in the search bar. This view contains all publication-specific information displayed in the variant view, and a list of all variants identified within the publication. Clicking on a variant ID or gene symbol allows navigation to the variant and gene views, respectively.

**Figure 1.**
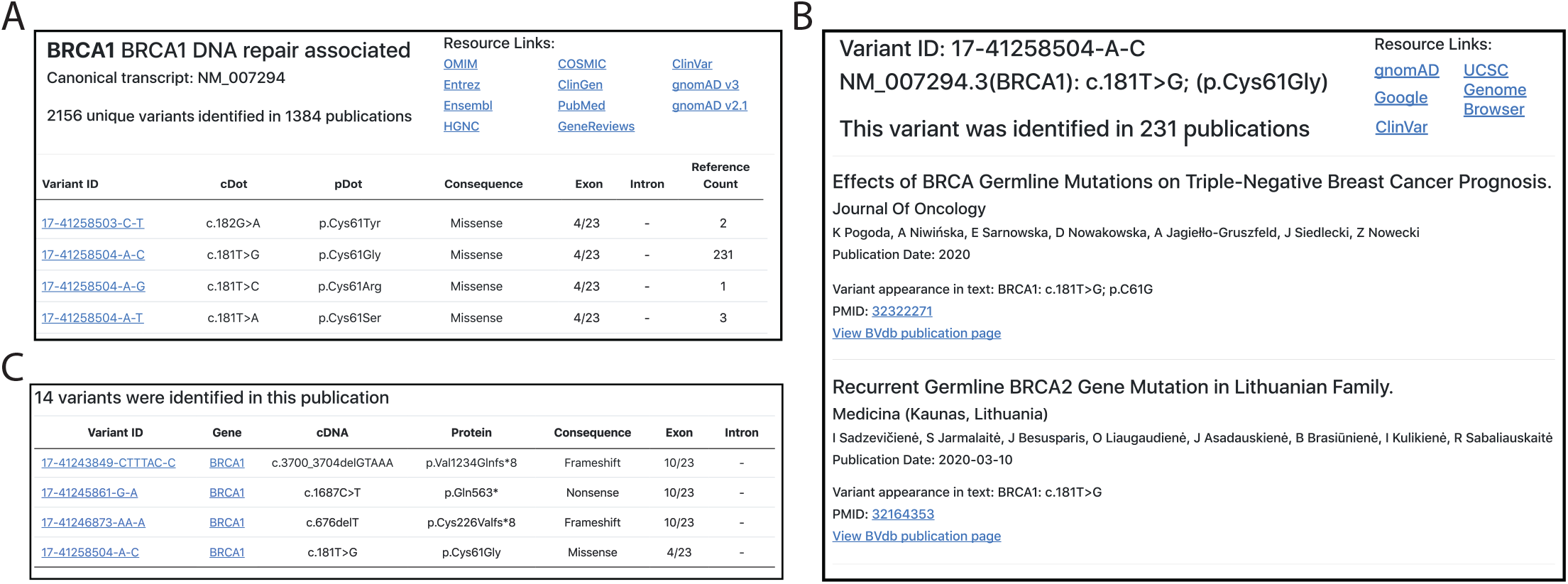
(A) The BVdb gene view, listing results from for the *BRCA1* gene symbol search term. (B) The variant view, displaying information and references describing the *BRCA1* c.181T>G; (p.Cys61Gly) variant. (C). The publication view, listing a subset of all variants identified in (Pogoda et al. 2020).

Ideal use-cases of the BVdb include variant interpretation, curation, and systematic literature review in clinical and research contexts. Use of the BVdb to gather evidence can reduce the time required to identify publications describing a given variant, a necessary and often rate-limiting process in clinical laboratory variant interpretation workflows (Ravichandran et al. 2019). Additionally, use of the BVdb can promote variant classification harmonization by ensuring all users have access to the same open access literature during variant interpretation (Garber et al. 2016). Furthermore, the BVdb can also facilitate variant reanalysis, by allowing easy identification of newly published literature describing previously evaluated variants (Baker et al. 2019).

## CONCLUSION

The BVdb is purpose-built to facilitate variant interpretation and curation in clinical and research contexts. The BVdb is a powerful tool that enables rapid lookup of primary literature describing genetic variation. Finally, automation ensures the BVdb can keep pace with the increasing volume of published literature and can add references describing genetic variants shortly after they become available.

